# Carry-over effects and plasticity to temperature shape phenology across life stages and generations

**DOI:** 10.64898/2026.09.10.749909

**Authors:** Rona Learmonth, Lea Beaupere, Ella F. Cole, Andrea Estandía, Ben C. Sheldon

## Abstract

Climate change is advancing spring phenology in temperate systems, with the potential to disrupt synchrony between trophic levels. Predicting these shifts requires understanding not only direct plastic responses to temperature, but also how plasticity at one stage carries over to shape timing at subsequent stages. We experimentally quantified direct thermal plasticity in phenology and its carry-over effects across the full life cycle of the winter moth (Operophtera brumata), a holometabolous insect whose fitness relies on synchrony with host plant budburst. Using a fully factorial split-clutch rearing experiment across four temperature treatments, we exposed individuals to contrasting conditions at each life stage and used structural equation modelling to partition direct and carry-over effects on phenology. Each life stage showed distinct plastic responses to temperature. Carry-over effects transmitted approximately 0.38 days/day of plastic advance to the next life stage on average, with the remainder absorbed by compensatory changes in the duration of the subsequent life stage. Together, these results show that carry-over effects propagate plastic responses across the life cycle, which are partially buffered by compensatory changes in developmental duration. Accurate predictions of phenological shifts under climate change therefore require models that account for carry-over effects and developmental compensation across life stages.

## Introduction

Rapid climate change is increasing temperatures globally (Calvin et al., 2023). Advances in spring phenology, particularly in temperate taxa, are among the most consistently reported ecological responses to this climate warming (Parmesan & Yohe, 2003; Renner & Zohner, 2018). Where interacting taxa have different temperature sensitivity, unequal advances can lead to trophic asynchrony (Samplonius et al., 2020; Thackeray et al., 2010), with negative fitness consequences for higher trophic levels (van Dis et al., 2023). Understanding the drivers of phenological shifts is key to predicting future asynchrony, as well as mitigating its fitness consequences.

Many documented phenological shifts have occurred through phenotypic plasticity to warming spring temperatures (Bonamour et al., 2019; Charmantier et al., 2008). Such phenotypic plasticity allows short-term adjustments in timing, which may promote or inhibit long-term adaptive evolutionary change (Martin et al., 2023). Understanding the extent and limits of phenological plasticity to temperature is therefore key to predicting future phenological changes and mismatch. However, a current focus on spring conditions may neglect the role of temperature changes across the rest of the year, especially given that rates of warming may be unequal between seasons (Cohen et al., 2012). There have been calls to consider how plasticity in timing across the whole year – not just spring – affect phenology and trophic asynchrony (Briscoe et al., 2012; Stålhandske et al., 2015, 2017). Failing to account for temperature responses across the entire year or life cycle can lead to incorrect predictions of spring phenology. For example, consideration of winter conditions revealed delays in butterfly emergence due to warmer winters (Stålhandske et al., 2017), and delays in arctic flowering due to greater winter snowfall (Bjorkman et al., 2015) despite warming springs. Developing our understanding of how phenology at one stage in the life cycle is shaped by cumulative developmental responses across the full annual cycle can thus improve our predictions of phenological shifts.

In taxa with life cycles involving metamorphosis between discrete life stages, phenology can be shaped across the life cycle by (i) plastic responses at each life stage, and (ii) carry-over effects between life stages. Each life stage may respond plastically to the conditions it experiences, but plastic responses to temperature may also interact across multiple life stages (Gray, 2013; Stillwell & Fox, 2005). Additionally, different life stages may vary in the magnitude or direction of their responses to the same environmental conditions (Briscoe et al., 2012; Marshall et al., 2020; Stålhandske et al., 2017; Weaving et al., 2022). Such differences may arise where life stages vary in their habitat, activity patterns, or temperature variation experienced (reviewed by Pottier et al., 2026). Carry-over effects, where an individual’s performance depends on its experience in a previous life stage (O’Connor et al., 2014), may propagate these plastic phenological shifts across life stages. Carry-over effects can be deleterious where they drive mismatch with environmental conditions or resource availability (Salis et al., 2018). Such phenological carry-over effects may be partially moderated by compensatory changes in subsequent life stages (e.g. Burraco et al., 2021; Orizaola et al., 2016), but the limits and fitness implications of such compensation remain uncertain. Therefore, as the timing of each life stage transition may respond to temperature differently, or be constrained by the responses of preceding stages, accurate predictions of phenology and mismatch must consider both stage-specific plasticity and carry-over effects across the full life cycle (Bonamour et al., 2019).

Winter moths (*Operophtera brumata*) are an ideal study system to understand how plasticity and carry-over effects shape phenology. Winter moth larvae are an economically and ecologically important defoliator of vegetation (Jepsen et al., 2013) and a food resource for other organisms, such as breeding birds (Coomes et al., 2025), so shifts in their phenology may have consequences across multiple trophic levels. Additionally, winter moths’ phenology is important for their own fitness, as synchrony between timing of hatching and host budburst determines their access to food resources (van Dis et al., 2023). Previous experimental studies of thermal plasticity in complex life cycles have used study systems such as *Drosophila melanogaster* (Klepsatel et al., 2023) or *Culex pipiens* mosquitoes (Gray, 2013), which have multiple generations each year, and can be exposed to different seasonal conditions at each generation. In contrast, winter moths are univoltine, with each non-overlapping life stage exposed to different seasonal conditions. Because winter moths are protandrous, with males emerging from pupation earlier than females (Topp & Kirsten, 1991), carry-over effects may further influence mating synchrony and reproductive success. This presents an ideal system in which to test how thermal conditions experienced across different seasons interact to shape life-cycle phenology.

Here, we investigate how thermal plasticity and carry-over effects across the entire life cycle of the winter moth shape its phenology using a fully factorial split-clutch rearing experiment across five temperature treatments (Fig. 1). To our knowledge, no previous study has factorially manipulated temperature across an entire life cycle to quantify both stage-specific plasticity and cross-stage carry-over effects. First, we quantify (i) the extent of plasticity in the timing of each life stage transition, and (ii) whether phenological shifts are carried over to subsequent life stages. Secondly, we assess how plastic phenological shifts affect the duration of each life stage. In line with previous studies, we predict plastic advances in the egg hatch and pupation timing with increasing temperature (Buse & Good, 1996; Salis et al., 2016; van Asch et al., 2007; van Dis et al., 2024), while adult emergence will be fastest at intermediate temperatures (Holliday, 1985; Peterson & Nilssen, 1998; Rattigan et al., 2026; Topp & Kirsten, 1991). We predict that, as previously reported, egg hatch timing will advance with the timing of adult emergence or oviposition (Learmonth et al., 2026; Rattigan et al., 2026; van Asch et al., 2010; Van Dongen et al., 1997). However, carry-over effects between other life stage transitions are uncertain, with mixed evidence for whether temperature-mediated timing at the larval phase influences subsequent life stages, or if compensatory changes in life stage duration are sufficient to overcome any carry-over effects (Salis et al., 2018; Senior et al., 2021; Topp & Kirsten, 1991). Understanding how temperature conditions across the entire year shape phenological responses across life stages is a key step towards building predictive models of spring phenology and mismatch

**Figure 1.**
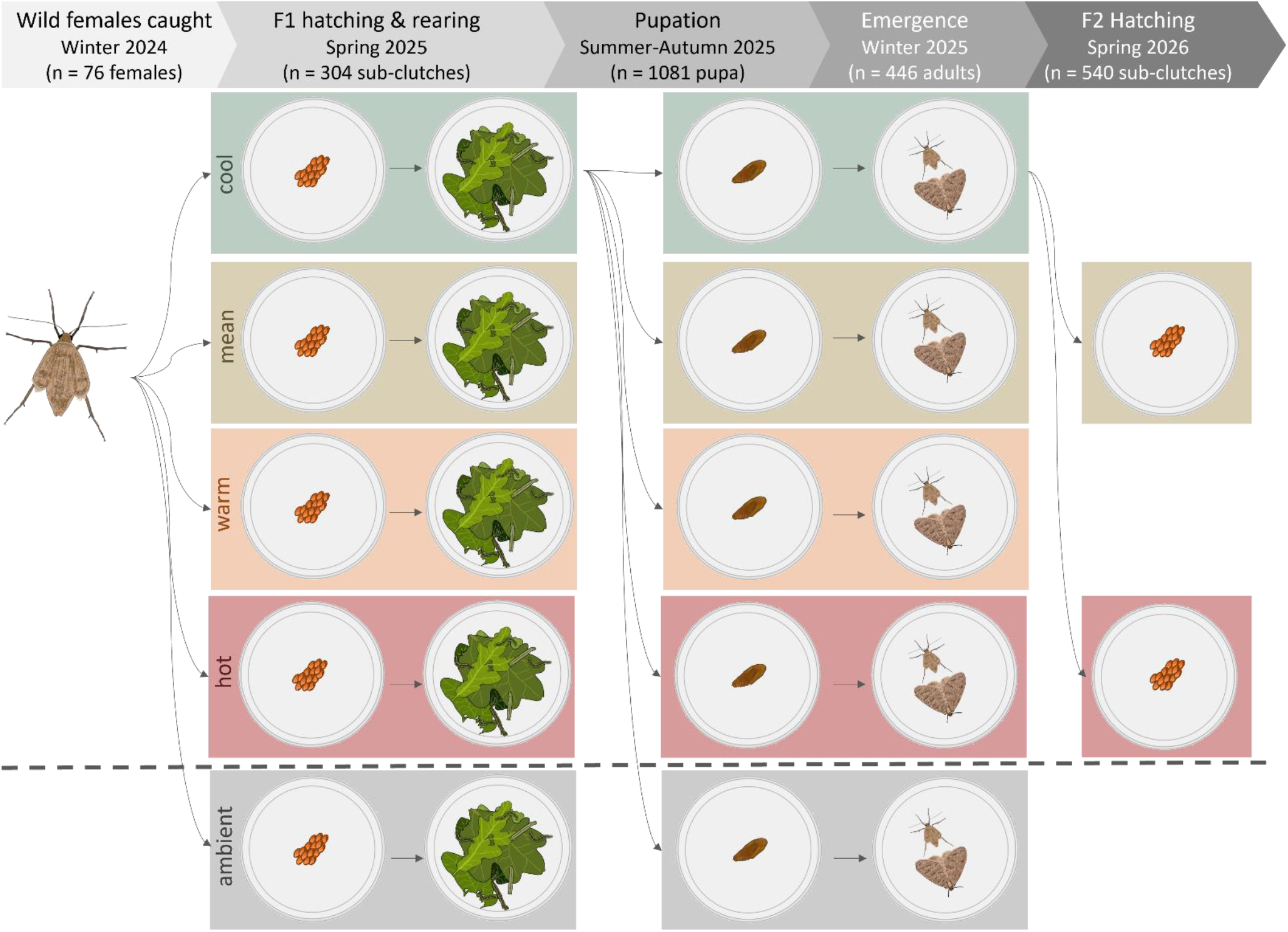
Schematic of cross-temperature rearing experiment. For a single female’s clutch, eggs are split into sub-clutches in hot, warm, mean, cool and ambient treatments (labelled and illustrated by box colour) for hatching and rearing to pupation. The timing of individual pupation is recorded, and each pupa reassigned across the temperature treatments. This reassignment is shown for the cool treatment only. After adult emergence, individuals were mated within matching early- and late-life treatment combinations, and their clutches re-split between mean and hot treatments only. Again, the reassignment is shown for the cool treatment only. Ambient treatments shown here below the dashed line were not used in full analyses as their temperature regimes are more variable and so not comparable to experimental temperatures (see Fig. S3). Sample sizes for each life stage are therefore given only for the four experimental treatments included in full analyses. For sample sizes including the ambient treatment, see Table S1.

## Methods

### Field sampling

In November-December 2024, we collected female winter moths from Wytham Woods, Oxford, (51°46′ N, 01°20′ W), a 385-ha area of mixed deciduous woodland. We installed paired ‘lobster pot’ geometrid moth traps (Jaworski & Sukovata, 2020, Fig. S1) on the north- and south-facing sides of 24 oaks (Quercus sp., see Fig. S2) across around 2 km of woodland. Sampled trees’ 2024 spring green-up dates, as calculated from curve fitting of drone-derived multispectral time series data (Klosterman & Richardson, 2017; Morley et al., 2025, SI Text), cover a normally distributed 29-day range. We collected females from each trap three times per week, storing each individually in a 15ml falcon tube with a roll of tissue paper for oviposition. 75% of all females caught laid eggs.

### Rearing experiment

#### Experimental temperature treatments

The eggs were stored outside until January, then split across four incubator (PHCbi MIR-154-PE) temperature treatments and one outdoor ambient treatment, which was kept in shade at John Krebs Field Station, Oxford (Fig. S3).

We used daily maximum, minimum, and mean from 50 years of temperature data for a 5 km grid square covering Wytham Woods (Met Office et al., 2024) to calculate a ‘mean’ treatment, which captures historical conditions as well as recent warming trends. To capture daily temperature patterns, we used the *chillr* package (Luedeling et al., 2023) to interpolate hourly temperatures from these daily measures, which were averaged across years. Due to limits in the number of programmes the incubators could store, we averaged temperature measures to two-hour intervals, resulting in a 12-step daily cycle, with each cycle repeated over two days. It should be noted that we had only one incubator per treatment, meaning treatment-level effects cannot be formally separated from incubator-specific effects. However, we found strikingly similar plastic responses to temperature regimes between the F1 and F2 generation, where treatment x incubator combinations were different, suggesting this pseudoreplication has limited impact on our results.

The contrasting temperature conditions were then derived by adding or subtracting standard deviations of monthly mean temperature from the same historical dataset. This produced the cool (- 1.5 SD), mean (historic average), warm (+1.5 SD), and hot (+2.5 SD) treatments.

Photoperiod has previously been shown to mediate carry-over effects in winter moths (Salis et al., 2018). Under climate change, we expect temperatures to increase but photoperiod to remain unchanged (Calvin et al., 2023), so the phenological response to temperature mediates the photoperiod experienced (i.e. early-hatching individuals will experience early-season photoperiod). Therefore, in our experiment we held the calendar-day photoperiod constant across treatments, so the photoperiod experienced by each individual would depend on their hatch timing. In the incubators, we applied a monthly mean photoperiod at two-hour resolution, calculated from mean sunrise and sunset times for the latitude of Wytham Woods.

#### Hatching and larval rearing

Each clutch with a minimum of 30 eggs (n = 76) was split into five sub-clutches with 5-15 eggs, stored in 60 ml plastic pots. We monitored the timing of egg hatching weekly from mid-February 2025. Winter moth eggs change colour from orange to blue-black around five days before emergence (Buse & Good, 1996; Embree, 1965). From first egg-darkening, we switched to daily checks of all clutches and added a cherry or oak leaf to the pot to prevent any period of starvation on emergence. These individuals and hatch monitoring methods are the same used in Learmonth et al. (2026).

We fed all caterpillars an excess of fresh leaves collected from trees near John Krebs Field Station, Oxford. At minimum every two days, we added new leaves and removed old leaves and frass. All winter moths were collected from oak trees, so we fed larvae on oak whenever possible. However, hatching began in the hot and ambient treatments before oak budburst, so larvae were initially fed on *Prunus* leaves until the first oak budburst in mid-April 2025. From this point onwards, all larvae were fed on an excess of fresh oak leaves. Therefore, we note that feeding regime has a confounding effect on response to temperature. This approach to feeding regime was taken to allow all larvae to be fed on newly-emerged leaves, as these are associated with greatest survival (van Dis et al., 2023). In a previous rearing experiment, survival probability and fitness have been shown to be similar for larvae fed on *Quercus robur* and *Prunus avium* (Weir, 2024), suggesting this feeding difference is unlikely to fully explain observed between-treatment differences. Nevertheless, we caution that the observed carry-over effects between life stages may be partly influenced by differences in feeding regime, rather than solely driven by temperature-mediated phenology.

We recorded larval weight to control for any temperature-dependent size differences. We first weighed larvae three weeks after the first hatching in that sub-clutch, as before this point larvae were too small for the 1 mg precision of the analytical balance (Ohaus PR124M). We re-weighed up to three randomly selected larvae from each sub-clutch at seven-day intervals until all larvae had pupated, using the mass/number of individuals as a measure of mean mass at each timepoint. We took mean larval weight of the sub-clutch at 35 days after first hatch as a proxy for larvae size, as this was the modal timepoint at which larval weight was maximised across treatments and clutches.

#### Pupation to adult emergence

We checked larvae for pupation daily. On the day of pupation, each individual pupa was transferred to a clean 60 ml plastic pot containing a cotton pad wetted with five drops of distilled water and approximately 25 ml of sterile vermiculite. Pupae were kept within their original temperature treatment until all larvae had pupated and were then reassigned across the five temperature treatments on the same day. Treatments were assigned randomly across the five treatments within each sub-clutch to maximise the number of families which were represented across all treatment combinations.

We monitored all pupae for adult emergence weekly from October, then every second day from first adult emergence on 3^rd^ November 2025. To balance the risks of desiccation or mould, we sprayed each pupa with distilled water every 16 days throughout pupation. Any pupae which became severely mouldy or desiccated were removed and assumed dead. Following the final adult emergence on 28^th^January 2026, we continued to monitor emergence every two days to ensure late-emerging individuals were sampled. In early March, we assumed that the remaining un-emerged pupae were unlikely to successfully emerge; we dissected all un-emerged pupae and found that none still appeared viable.

On adult emergence, we recorded date, sex, and female weight, which correlates with fitness in terms of number of eggs laid (Fig S4). We did not record male weight, as the winged males are more difficult to weigh and the effect of male size on female fitness is unclear (Van Dongen et al., 1999). Adult moths were mated to an unrelated individual within their treatment combination, such that they had experienced the same egg and pupal temperature treatments. We prioritised pairing older adults to maximise the number of individuals able to mate. Each mated pair was stored in a clean 60 ml plastic pot with two rounds of tissue paper on the base for oviposition. Each adult mated with only one individual, and pairs were stored together until the female’s death, after which the adults were removed. Adults were stored in the same temperature treatment as for pupation.

#### Hatching F2 generation

Clutches of eggs were kept in their parental treatment until January. We counted the number of eggs in each clutch, and split each clutch with at least 30 fertilised orange eggs (Embree, 1965) between the mean and hot treatments in two sub-clutches of 15 eggs. All eggs were transferred into their new treatment on the same day, which occurred before the onset of significant temperature sensitivity in the eggs (Van Dis et al., 2021).

We monitored the sub-clutches for egg hatching every second day from late February, recording hatch time of each larva. We continued to monitor hatching until three weeks after the last hatching event, at which point unhatched eggs were assumed inviable.

## Statistical analysis

### Data processing

We conducted all analyses in R (R Core Team, 2025). As in Learmonth et al. (2026), we calculated half-hatch day for each clutch as a measure of mean hatch phenology for a sub-clutch. Half-hatch day represents the point at which 50% of eggs had hatched, calculated from the proportion of eggs hatched by experimental day for each sub-clutch in each treatment corrected for deaths during development. Pupation and adult emergence were recorded directly at the individual level. Half-hatch day for eggs hatching in spring 2026 was extracted as previously (Learmonth et al., 2026) but used the raw number of individuals hatched per day, as we did not rear any larvae and so did not need to account for new births and deaths in the same way.

We converted categorical treatment factors into a pseudo-continuous variable to allow direct interpretation of how phenology shifts per degree temperature change. We used a standardised time period across all treatments, as using time windows relative to each treatment’s phenological timing would introduce circularity in models of the impacts of temperature on phenology. Therefore, we used mean temperature of the entire life phase up to the median date of life stage transition across all treatments as a pseudo-continuous temperature measure for each stage. For the egg phase, this lasted from when the eggs entered the incubators until median half-hatch day; for larvae, from median half-hatch day until median pupation date, and for adults from the date pupae were assigned across incubators to median adult emergence time. Because all treatments were generated based on standard deviations from the mean, z-scaled temperature differences between treatments remain very similar (within 0.001) regardless of the time period selected.

We compared results from the experimental treatments to the ambient treatment to confirm that the relationships under experimental conditions match expectations when exposed to natural variability (see SI Text, Fig. S5).

### Modelling: timing of life stages

We used a structural equation modelling (SEM) approach in *cmdstanr* (Gabry et al., 2025) to estimate the direct and indirect effects of plasticity and carry-over effects on subsequent life stages. Here, plasticity refers to direct phenological responses to temperature within a life stage, while carry-over effects refer to shifts in timing at one stage predicted by timing at a previous stage. The grand model was composed of four models of the predictors of (i) mean egg hatch timing at the sub-clutch level (*half-hatch day*), (ii) individual timing of pupation, (iii) individual timing of adult emergence, and (iv) mean egg hatch timing of F2 sub-clutches. In each case, all predictor variables were z-scaled.

We modelled how mean egg hatch timing at the sub-clutch level (half-hatch day) was predicted by the mother’s catch date as a proxy for timing of adult emergence, the mean temperature of the egg phase, and mother’s identity as a group-level effect. We then modelled how individual timing of pupation was predicted indirectly by these factors via the carry-over effect of hatch timing, as well as directly by mean temperature of the larval phase, larval mass at 35 days, and mother’s identity as a group-level effect. We modelled how individual timing of adult emergence was predicted indirectly by all these paths via carry-over of pupation timing, as well as directly by mean temperature of the pupal phase, larval weight at 35 days, and female identity as a group-level effect. We also included a quadratic effect of pupal temperature on adult emergence based on previous findings that adult emergence is earliest at intermediate temperatures (Peterson & Nilssen, 1998; Rattigan et al., 2026; Topp & Kirsten, 1991). For F2 hatch timing, we modelled the effects of carry-over from maternal emergence time, temperature experienced, and group-level effects of grandmother and mother identity.

Each model was run using a Gaussian distribution on six Markov chains, with 10,000 iterations each, at an adapt delta of 0.95 and maximum tree depth of 12. We carried out posterior predictive checks, comparing 1000 draws of the posterior predictive distribution with the observed data, which confirmed that the model reflects real patterns in the data. We also checked model diagnostics to confirm the chains were appropriately mixed, and 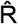 values did not diverge from 1.

### Modelling: duration of life stages

While modelling the timing of each life stage transition captures the shifts in phenological events, it does not consider how these shifts affect the organism’s individual life history. Consequently, we also modelled how plasticity and carry-over effects determine the duration of each life stage to explore whether changes in phenology drive compensatory responses. We note that compensatory changes in life stage duration can also be inferred from the timing of life stages model as the magnitude of phenological shift not transmitted to the next stage (1 – carry-over effect). However, by directly modelling the duration of each life stage we can explicitly link developmental rates between life stages and environmental contexts.

We first calculated the duration of each life stage at the relevant scales of organisation. The duration of the egg stage was taken as the time from the mother’s capture date to the sub-clutch half-hatch day. This measure includes some error, as oviposition time was not recorded directly, but is assumed to occur within around four days of adult emergence at ambient temperatures (Rattigan et al., 2026). This four-day error measurement would represent 2.92% of the egg phase (given average duration of 136.96 (± 0.87) days). Duration of the larval phase was calculated as the time from sub-clutch half-hatch day to individual pupation time, though we note that early-hatching eggs within a clutch may also have pupated earlier. The duration of the pupal phase was calculated as time from pupation to adult emergence. We did not assess the time taken from adult emergence to death, as the duration of survival after mating and oviposition should not affect the subsequent generation.

We then used a SEM in *cmdstanr* (Gabry et al., 2025) with similar structure to the one used to investigate timing of phenological events to examine how duration of life stages was impacted by plasticity and carry-over effects. The grand model was composed of three sub models. First, the duration of the egg phase was predicted by the temperature it experienced. Second, the duration of the larval phase was predicted by the temperature experienced, a carry-over effect of the egg duration, and the mean larval weight at 35 days for the sub-clutch. Finally, pupa duration was predicted by larval duration, temperature as a linear and quadratic term, and larval weight. The quadratic effect of temperature was included because, if pupal emergence is earliest at intermediate temperatures (Topp & Kirsten, 1991), pupation was expected to terminate earlier and therefore be shorter at intermediate relative to extreme temperatures. The mother’s identity was included as a group-level effect in each model. As previously, all variables were z-scaled before inclusion in modelling, and the model was run using a Gaussian distribution on six chains for 10,000 iterations each at an adapt delta of 0.95 and maximum tree depth of 12. We performed posterior predictive checks to confirm the models reflect real patterns in the data and checked model diagnostics to confirm mixing of chains and 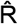 values of 1.00.

### Effect of temperature on protandry

Winter moths are protandrous, with males emerging several days before females to maximise mating opportunities (Topp & Kirsten, 1991). Thus, to examine how mating opportunities are potentially impacted by responses to temperature, we investigated how sex impacts emergence time across treatments. We modelled how emergence time was predicted by sex, and its interactions with early-life and late-life treatment in the package *brms* (Bürkner et al., 2024). We included mother’s identity as a group-level factor. The model was run using a gaussian distribution on four Markov chains, with 5000 iterations each at an adapt delta of 0.95 and maximum tree depth of 12. We carried out posterior predictive checks and confirmed model diagnostics were appropriate

## Results

### Plasticity to temperature

Structural equation modelling (SEM) of the impacts of plasticity and carry-over effects on the timing of life stage transitions demonstrated plastic responses to temperature at all life stages (Fig. 2). Timing of egg hatch advanced with increasing temperature equally in the F1 and F2 generations (F1: estimate = -0.95, CI = [-0.97, -0.92], shown in Fig 2; F2: estimate = -0.96, CI = [-0.98, -0.94]). The timing of pupation likewise advanced with increasing temperatures (estimate = -1.01, CI = [-1.13, -0.89]), while adult emergence had a quadratic relationship with temperature (linear estimate: -0.26, CI = [-0.33, -0.19], quadratic estimate = 0.23, CI = [0.15, 0.31]), such that emergence was earliest at intermediate temperatures.

**Figure 2.**
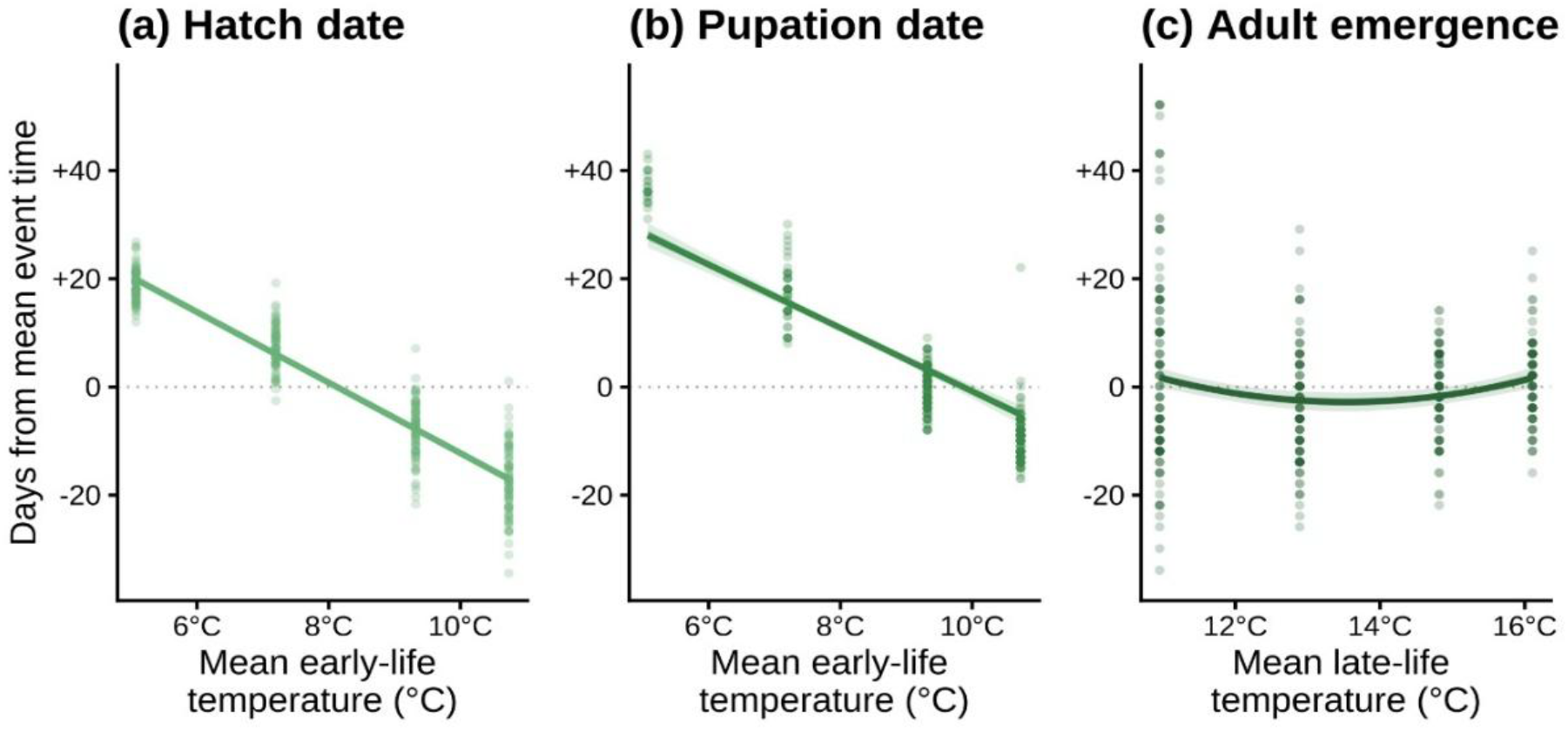
Plasticity across life stages. Conditional effects plot summarising the degree of temperature plasticity in (**a**) hatch, (**b**) pupation, and (**c**) adult emergence timing for the F1 generation only. The y-axis shows the number of days divergence from the mean, to make estimates comparable across stages, which is plotted against mean temperature experienced over the course of that life stage. In each case, the dotted line at zero shows the mean phenology of the life stage. The coloured line indicates the model predictions, once carry-over effects, larval weight, and group-level effects of mother identity are controlled, surrounded by its 90% credible intervals. The points represent the underlying raw data, where transparency of colour indicates number of points overlaid.

### Plasticity & carry-over effects shape phenology of life stage transitions

Carry-over effects occurred between all life stages we examined, such that a delay in one life stage transition was associated with a delay in the next life stage transition (Fig. 3). This means that a larva which hatched late as an egg would pupate late, regardless of the temperature conditions the larva experienced. The strength of this phenological carry-over effect varied somewhat between life stages. The strongest carry-over effect was of the timing of pupation on adult emergence (estimate = 0.48, CI = [0.42, 0.54]), followed by timing of hatch on pupation (estimate = 0.34, CI = [0.21, 0.45]), and timing of adult emergence on F2 hatching (estimate = 0.31, CI = [0.27, 0.35]). Across the full F1 carry-over pathway (hatch **→** pupation **→** adult emergence), for each day delay in egg hatching, adult emergence was delayed by 0.12 days (CI = [0.08, 0.17]). Across the full experiment, direct plasticity to temperature explained 73.4% (CI = [23.2%, 29.8%]) of variation in timing, while carry-over explained 26.6% (CI = [23.2%, 29.8%]).

**Figure 3.**
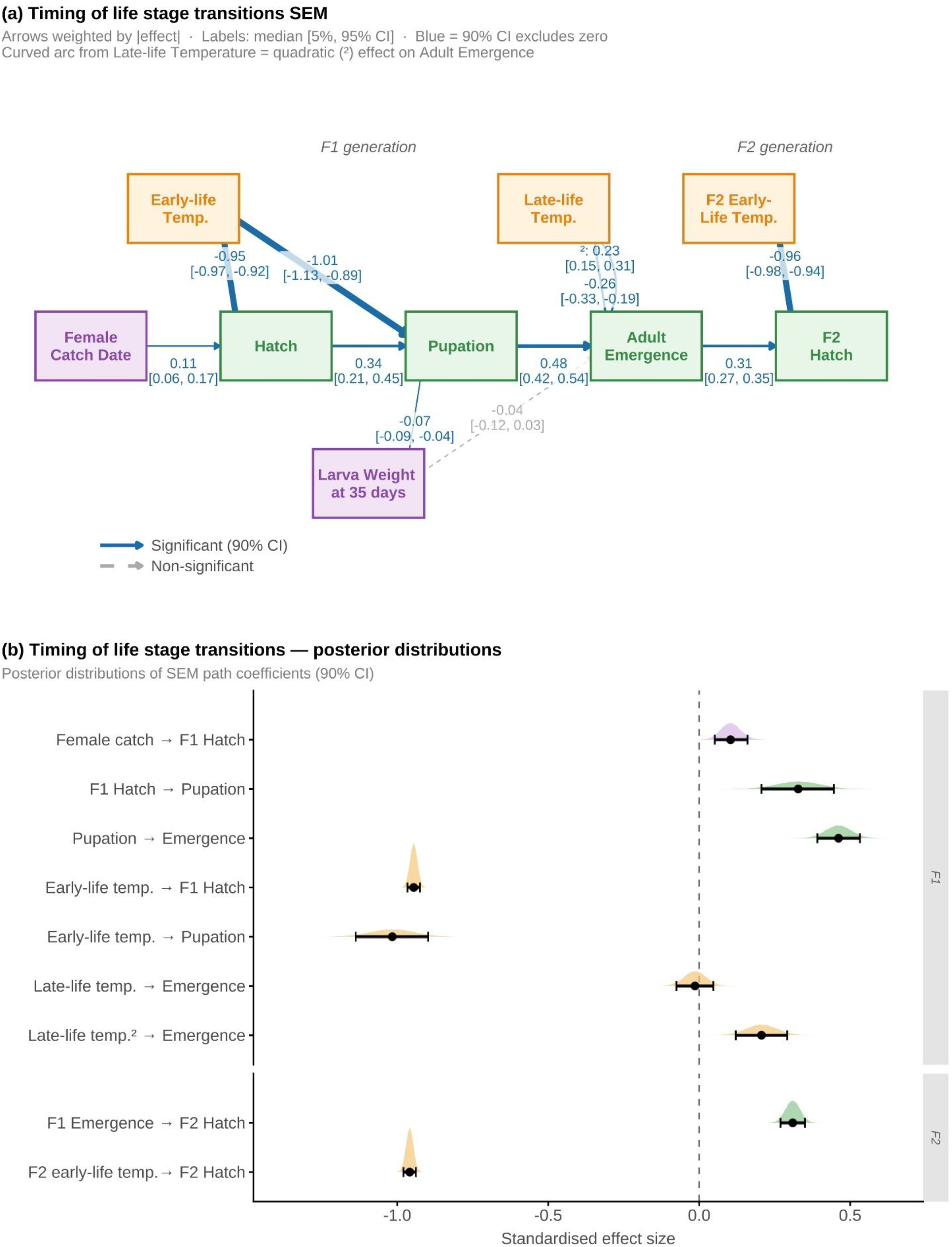
Timing of life stage transition SEM. (**a**) Posterior path coefficients showing the relationship between covariates (purple), timing of life stages (green) and temperature treatments experienced (yellow) for F1 and F2 individuals. In each case, the size of the arrow indicates the median effect size, which is further printed, with its credible intervals, beside the arrow. All data were z-scaled prior to modelling. Both the linear (lower) and quadratic path (upper) for the effect of late-life temperature on adult emergence time are shown. (**b**) Model posterior distributions for the standardised effects of treatment (temperature plasticity) and carry-over effects on each life stage transition.

The timing of F1 hatching also advanced with the capture date of the mother, but this was treated as a covariate rather than a strict carry-over effect. While the capture date of winter moths from the wild is often used as a proxy for their emergence timing (Learmonth et al., 2026; Van Dongen et al., 1997), it differs from all other measures used here as it was collected at a coarser time-scale of three checks per week, was not experimentally manipulated, and includes greater uncertainty about the actual timing of the life stage transition. Increases in larval weight (at 35 days, the modal timing of maximum weight) were associated with slightly earlier pupation (estimate = -0.07, CI = [-0.09, -0.04]), while there was no effect on timing of adult emergence.

### Plastic changes in duration of life stages

Following similar structure to the timing SEM, we modelled how the duration of each life stage was impacted by plasticity to temperature and the duration of the preceding life stage. As for timing of life stage transitions, we found that higher temperatures reduced the duration the egg and larval phase (egg: estimate = -0.90, CI = [-0.92, - 0.88]; larva: estimate = -2.66, CI = [-2.95, -2.38], Fig. 4), with a positive quadratic effect of temperature on pupa duration (estimate = 0.18, CI = [0.08, 0.27]).

**Figure 4.**
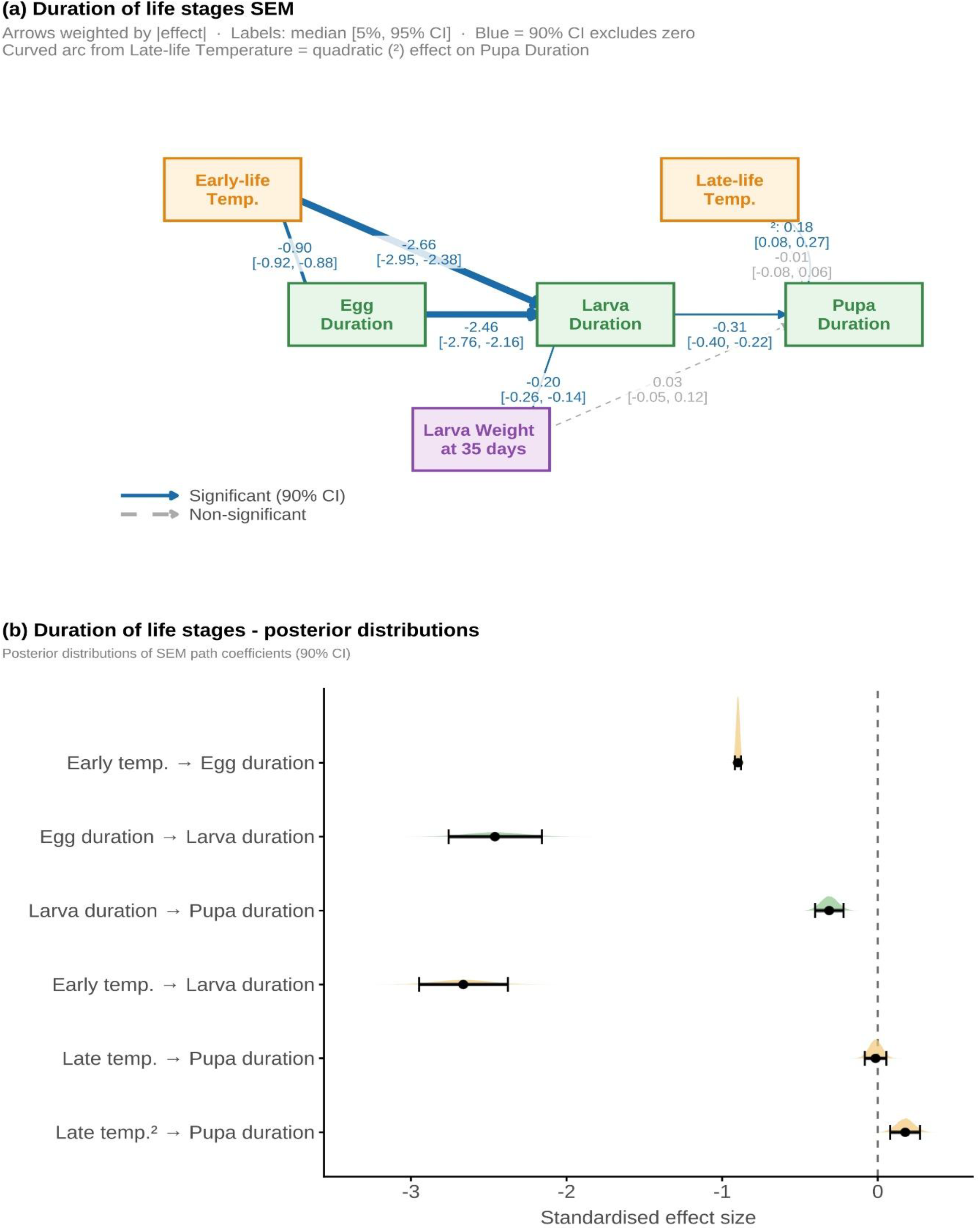
Duration of life stage transition SEM. (**a**) Posterior path coefficients showing the relationship between covariates (purple), timing of life stages (green) and temperature treatments experienced (yellow). In each case, the size of the arrow indicates the median effect size, which is further printed, with its credible intervals, beside the arrow. All data were z-scaled prior to modelling. (**b**) Standardised effects of treatment (temperature plasticity) and carry-over effects on the duration of each life stage.

The duration of each life stage decreased as the duration of the previous life stage increased (larva: estimate = -2.46, CI = [-2.76, -2.16]; pupa: estimate = -0.31, CI = [-0.40, -0.22], Fig. 4). In other words, a longer egg phase was followed by a shorter larval phase, or a long larval phase followed by a shorter pupal phase. As previously, greater larval weight at 35 days was associated with a shorter larval phase (estimate = -0.20, CI = [-0.26, -0.140) but no impact on the duration of the pupal phase (estimate = 0.03, CI = [-0.05, 0.12]).

### Temperature influences the extent of protandry

We modelled how temperature treatments at multiple life stages interact with the relative timing of emergence of males and females. For both sexes, emergence became earlier with increasing early-life temperature, with hot early-life treatment individuals emerging 11.52

[-15.41, -7.69] days earlier than individuals from the mean early-life treatment. For both sexes, mean late-life (i.e. pupal) temperature was associated with earlier emergence (mean: estimate = 63.79 [59.61, 67.88]) than any other treatment. Family identity explained a significant portion of variation, with among-mother differences of 5.70 days [4.34, 7.29].

The temperature experienced at each life stage influenced the extent to which males emerged earlier than females. For individuals exposed to hot, warm or mean temperature conditions in early- and/or late-life, males emerged slightly earlier than females, though with considerable overlap (Fig. 5).

**Figure 5.**
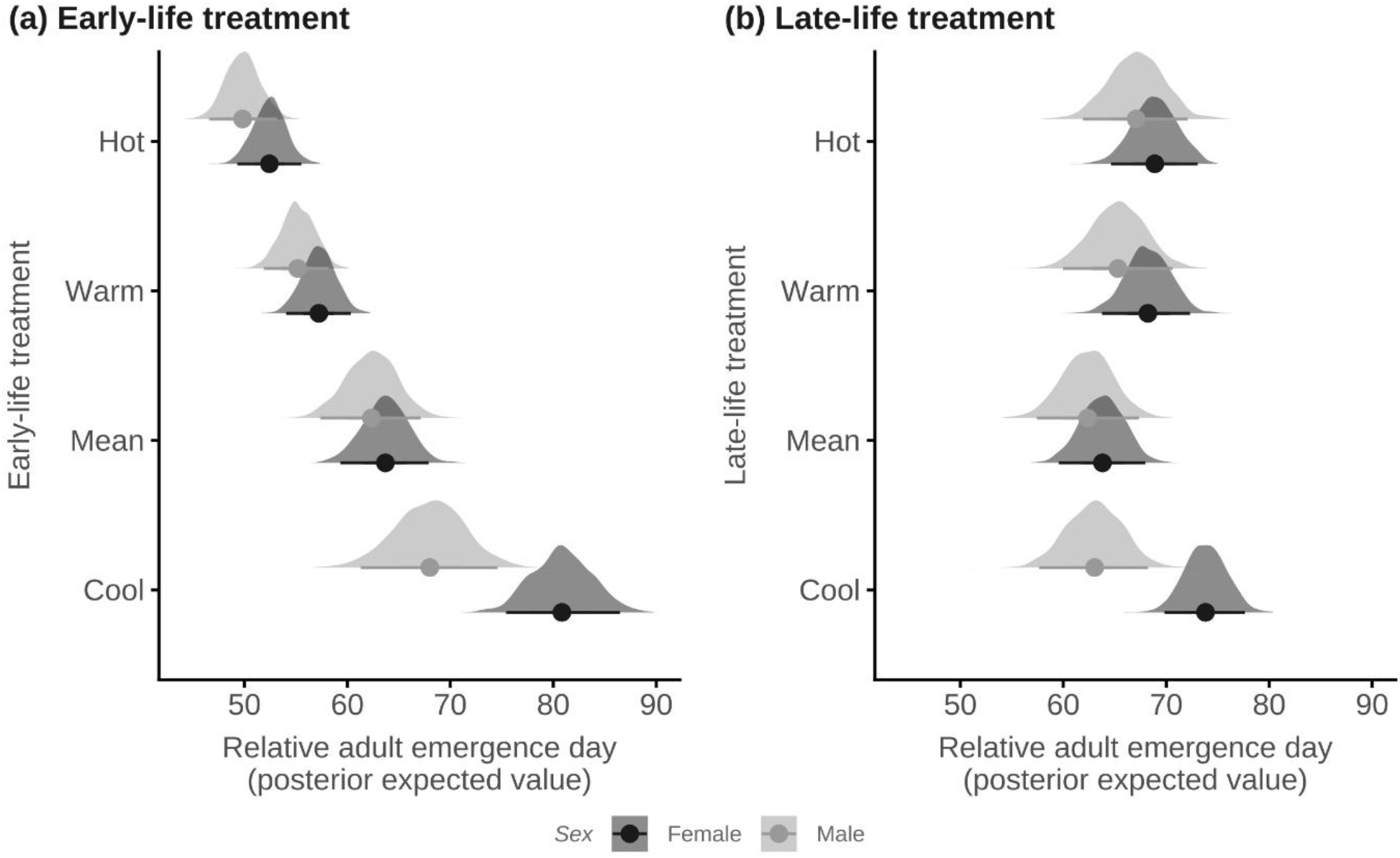
Temperature drives changes in the extent of protandry (male-first adult emergence). The plots show the posterior distributions of emergence timing for each sex at each (a) early life (egg and larva) and (b) late life (pupa and adult) temperature, where the effects of other temperature treatment and family identity are controlled. In each case, males generally emerge first, but this difference is magnified in those individuals exposed to the cool treatment, either in early or late life.

At the reference condition (mean early- and mean late-life treatment), males emerged slightly earlier than females, though with a high level of overlap (estimate = - 1.40, [-7.80, 4.74]). However, in the cool late-life treatment, males emerged 10.92 [-22.17, -0.03] days earlier than females. Similarly, following exposure to the cool early-life treatment, males emerged 12.22 days [-31.20, 3.39] earlier than females, though this estimate was associated with higher uncertainty and the credible intervals overlap zero.

## Discussion

We experimentally manipulated temperature across two generations in winter moths to assess how temperature shapes phenology through plastic responses, carry-over effects, and compensatory developmental changes. We found plastic advances in the timing of every life stage, indicating that warming temperatures will generally accelerate timing across the life cycle. We further report phenological carry-over effects which could amplify phenological shifts under climate warming. However, we also found compensatory changes in the duration of each life stage given the duration of the previous one, which could mitigate the impacts of plasticity and carry-over effects on phenology. Together, our results indicate the complexity of phenological responses to temperature when considered across the life cycle.

We found advances in phenology with increasing temperature at hatching and pupation, while adult emergence was earliest at intermediate temperatures. These findings align with previous studies of winter moth egg hatching (Buse & Good, 1996; Salis et al., 2016; van Asch et al., 2007; Van Dis, 2023) and adult emergence (Peterson & Nilssen, 1998; Rattigan et al., 2026; Topp & Kirsten, 1991). However, to our knowledge this study is the first to directly compare thermal plasticity of phenology across the full life cycle. Our approach allowed us to quantify the relative thermal plasticity of phenology at each stage, revealing that temperature sensitivity differs across stages exposed to different seasonal environments.

While phenological plasticity to temperature can be beneficial, allowing rapid tracking of environmental conditions (Charmantier et al., 2008; López-Idiáquez et al., 2026), it can also be maladaptive. In the Netherlands, plastic advances in winter moth hatch timing historically exceeded advances in tree budburst, resulting in trophic mismatch (Visser & Holleman, 2001, though see van Asch et al., 2013). This was linked to strong selection on hatch timing, resulting in rapid adaptive evolution in phenology (van Asch et al., 2013). This highlights the importance of separating the roles of evolutionary and plastic change in climate change responses to predict how populations may cope or adapt to climate change (Gienapp et al., 2008). By quantifying phenological plasticity at each stage, we can advance understanding of how plasticity may affect selection and evolutionary responses (Gibert et al., 2019). A key next step is to examine how the non-additive effects of plasticity at each life stage alter synchrony with food plants to predict evolutionary responses.

As well as mediating trophic synchrony, we show that plasticity to temperature may also influence synchrony of emergence between males and females. The extent to which males emerged earlier than females (protandry) was impacted by the temperature treatment they experienced. Where moths were exposed to the ‘cool’ treatment, either in early-(egg, larva) or late-life (pupa, adult), males emerged 10-12 days earlier than females. Given the average lifespan of the males is only around 7-10 days in this study and others (Topp & Kirsten, 1991; Van Dongen et al., 1999), this increase in protandry could restrict mating success and potentially reduce population sizes. To our knowledge, temperature-dependence of winter moth protandry has not previously been demonstrated, but feeding on alternative plants has been shown to extend or eliminate protandry relative to the original host (Tikkanen et al., 2000). Additionally, in orange-tip butterflies (*Anthocharis cardamines*), there is experimental evidence that relative development time of males and females is temperature sensitive (Stålhandske et al., 2015). Given that temperature-mediated shifts in the extent of protandry could potentially restrict mating opportunities and so affect fitness at the population level, further work should address how protandry will shift under future climate scenarios, and its demographic consequences.

In our experiment, we found phenological carry-over effects between successive stages, where late timing at one stage was associated with later timing at the next. Carry-over effects from maternal emergence timing to egg hatch timing were expected based on previous work (Salis et al., 2018; Van Dongen et al., 1997). However, perhaps more surprising is the persistence of phenological carry-over effects from egg hatching to adult emergence. By multiplying across intermediate paths, we found that adult emergence advances 0.12 days (CI: [0.07, 0.17]) for every one-day advance in egg hatching.. While a previous study in winter moths reported a phenological carry-over effect of larval photoperiod on adult emergence, the authors stress that a carry-over effect of hatch timing on adult emergence would be deleterious (Salis et al., 2018). Our results demonstrate that phenological carry-over effects occur across the full life cycle and into the next generation.

Whether such carry-over effects are maladaptive depends on the extent to which temperature conditions at one life stage or generation predict those in the next (temporal autocorrelation). In Wytham Woods, our study site, temperatures during the late-life treatment window when adults emerge were only weakly correlated with temperatures in the following early-life treatment window, when their eggs hatch (r = 0.22 [-0.06, 0.47], see *SI Text*). Thus, temperature cues experienced at adult emergence do not reliably predict the conditions larvae will experience, meaning the adult-offspring carry-over effect may increase the risk of maladaptive shifts in hatch timing away from the optimum. The optimal timing of hatching varies between years (van Asch et al., 2013). Additionally, early-life conditions between years are only modestly autocorrelated (r = 0.33 [0.06, 0.56], see *SI Text*) so the continued carry-over effects on the following generations may then amplify phenological shifts. Together, these results suggest phenological carry-over effects across life stages and generations may lead to greater phenological shifts or more severe asynchrony than would be predicted based on the focal spring temperatures alone. Consequently, we emphasise the importance of considering phenological responses to temperature across the full life cycle when predicting phenological responses to climate change.

While carry-over effects can propagate across the life cycle, we observed partial compensation in the duration of each life stage. Where one stage was extended, the next was contracted: for example, where eggs in the cool treatment hatched late, they experienced a shorter larval phase. Other experimental studies of winter moths have previously demonstrated similar effects only in the pupal stage following larval development at warmer temperatures (20°C vs 12.5°C) or late-season photoperiod (Salis et al., 2018; Topp & Kirsten, 1991). This compensation is consistent with the idea that phenological carry-over effects can be mitigated by shifts in the duration of life stages (Salis et al., 2018). Such changes in the relative duration of each life stage may reduce the extent to which carry-over effects exacerbate phenological asynchrony or disrupt mating synchrony.

Phenological compensatory mechanisms have been studied frequently in anuran frogs, which accelerate their larval development in response to delayed hatching or breeding (Richter-Boix et al., 2014). While this compensatory rapid growth is adaptive and allows maintenance of synchrony with the environment, it is also linked to reduced development of antipredator defences (Orizaola et al., 2016). Additionally, compensation can be limited under harsh environmental conditions (Burraco et al., 2021). Taken together, this evidence from frogs demonstrates that compensatory mechanisms like those we report in the winter moth can be crucial to the maintenance of synchrony in complex life cycles. However, it also highlights the need for future work to assess whether this compensation is associated with fitness consequences. For example, reduced larval phases may alter growth curves, potentially reducing adult weight and clutch size. Further work should also assess the extent to which this compensation functions under harsh vs benign environments, for example in terms of resource availability or temperature conditions.

## Conclusion

To conclude, we demonstrate experimentally that temperature conditions across the annual cycle shape phenology, through both direct plastic responses and developmental carry-over effects. Although compensatory shifts in stage duration may partially buffer these effects, phenological responses can nevertheless propagate across life stages, with potential implications for phenological shifts and trophic synchrony. Our findings highlight the importance of integrating multi-stage plasticity and carry-over dynamics into predictive models of phenological change.

## Acknowledgements

We thank Dr David López-Idiáquez for help monitoring emergence from pupae, and Celestine Adelmant for the illustrations used in Figure 1.

## Author Contributions

Conceptualisation: RL, EFC, BCS; Data collection: RL, LB; Formal analysis: RL; Funding acquisition: BCS; Supervision: AE, EFC, BCS; Investigation: RL, LB; Methodology: RL, LB; Visualisation: RL, Writing – Original Draft: RL; Writing – review and editing: RL, LB, AE, EFC, BCS

