## Supplementary Information for "Carry-over effects and plasticity to temperature shape phenology across life stages and generations"

#### SI text:

##### Drone-derived green-up estimates

Drones were flown over Wytham Woods every three days, weather permitting, from March to May each year. The NDVI images from these flights were stitched into orthomosaics, from which tree crowns for the focal oaks were manually delineated. For each tree, mean daily NDVI values for the central 75% of the tree crown were extracted, and the date at which NDVI reached 50% in the green-up curve identified as a proxy for budburst date. These methods use the same dataset and protocol as described in Morley et al. (2025).

##### Treatment vs ambient comparison

The ambient treatment experienced much greater temperature variability than the incubators (Fig. S1). To test whether the experimental effects in incubators matched those expected under conditions of natural variability, we included an ambient outdoor treatment in the factorial design. We compared the plasticity reported for the experimental treatments to those seen in the ambient treatment (Fig. S2).

##### Autocorrelation of early- vs late-life treatment windows in Wytham Woods

Using the same 1965-2016 temperature dataset used to produce the experimental temperature treatments, we calculated the one year lagged correlation of the early-life (day of year 35-139) and late-life (day of year 181-337) periods. We calculated the between-year Pearson correlation for early and late-life periods respectively to determine how conditions in a particular life stage in year t predict those in year t+1. We further calculated how temperature in the late-life period in year t predicts temperature in the early-life period in year t+1 to reflect how conditions on adult emergence predict those experienced by their offspring the following spring.

### SI tables & figures

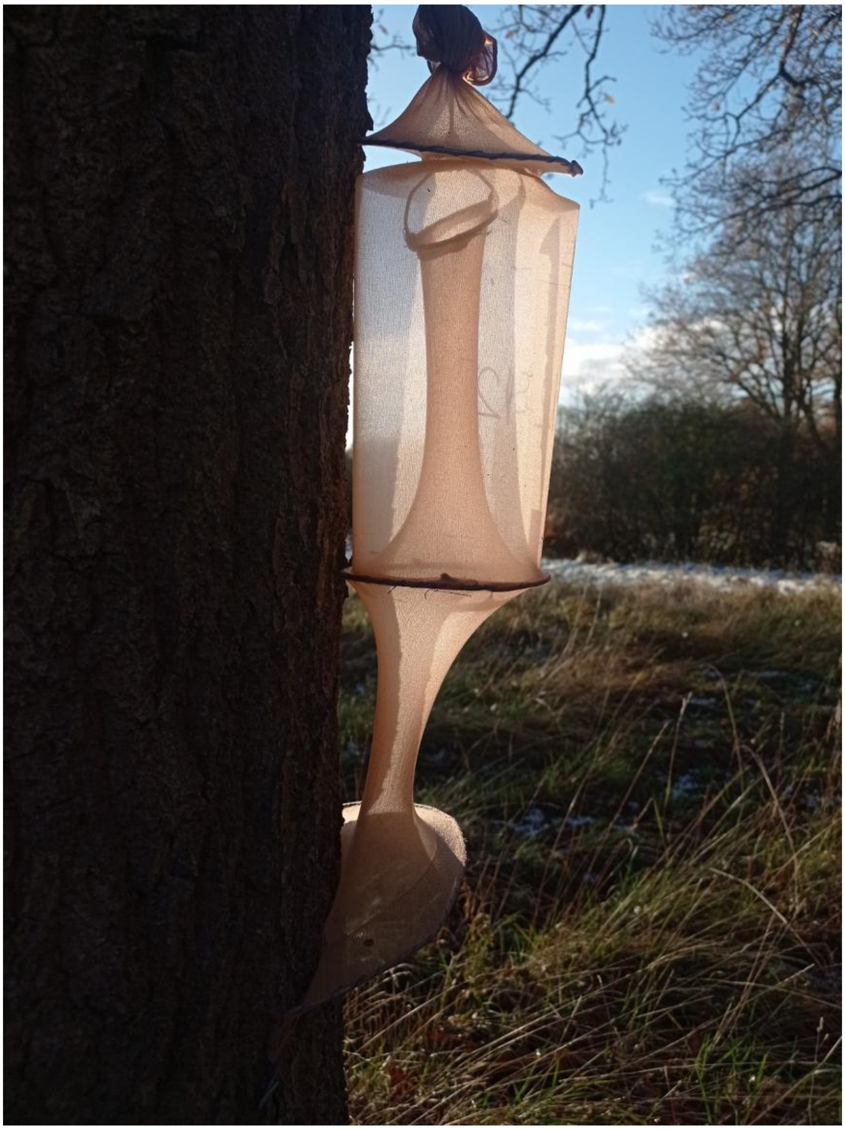

**Figure S1: Gradwell trap used to catch wild female winter moths in Wytham Woods.** The same traps were used in each iteration of trapping

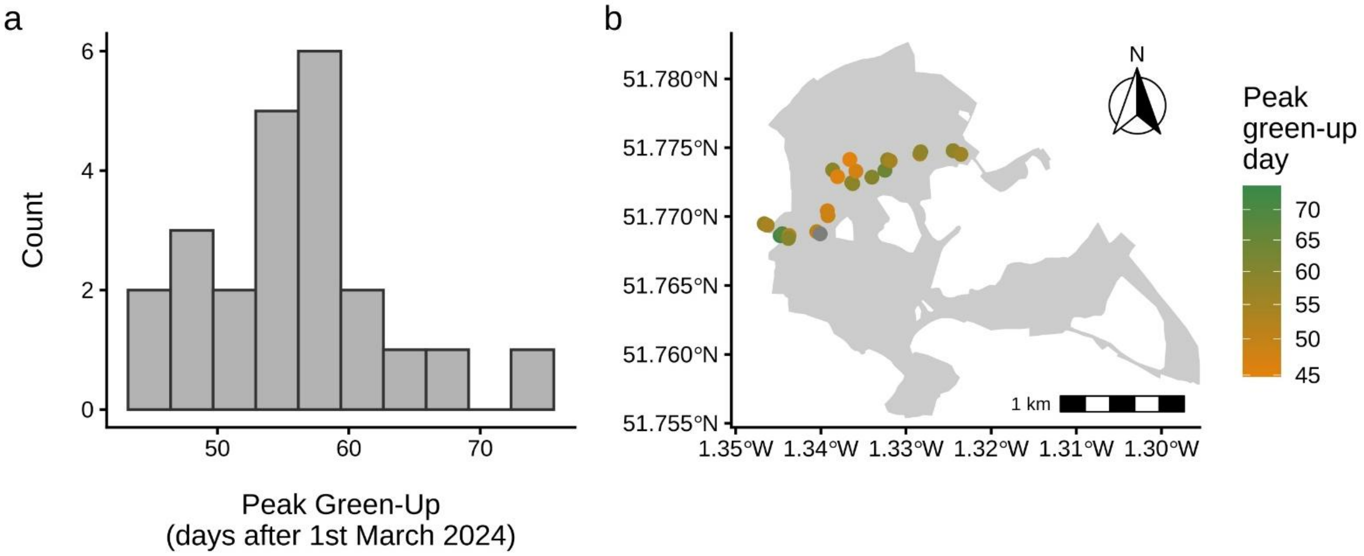


**Figure S2: Sampled trees phenology distribution and location.** (a) Histogram showing the distribution of 2024 spring green-up dates for trees sampled for winter moths. (b) Map of trees in Wytham Woods, UK, sampled for winter moths in 2024. The colour of each dot indicates its relative green-up date.

**Table S1:** Sample sizes for all experimental stages including ambient early/late-life treatments

| **Stage** | **Experimental treatments** | **Sample size** |
|---|---|---|
| **Female catch dates** | n/a | 75 females |
| **Egg hatching** | Hot, warm, mean, cool, ambient | 380 sub-clutches |
| **Pupation** | Hot, warm, mean, cool, ambient | 1566 individuals |
| **Adult emergence** | Hot, warm, mean, cool, ambient | 773 individuals |
| **F2 egg hatching** | Hot, mean | 578 sub-clutches |

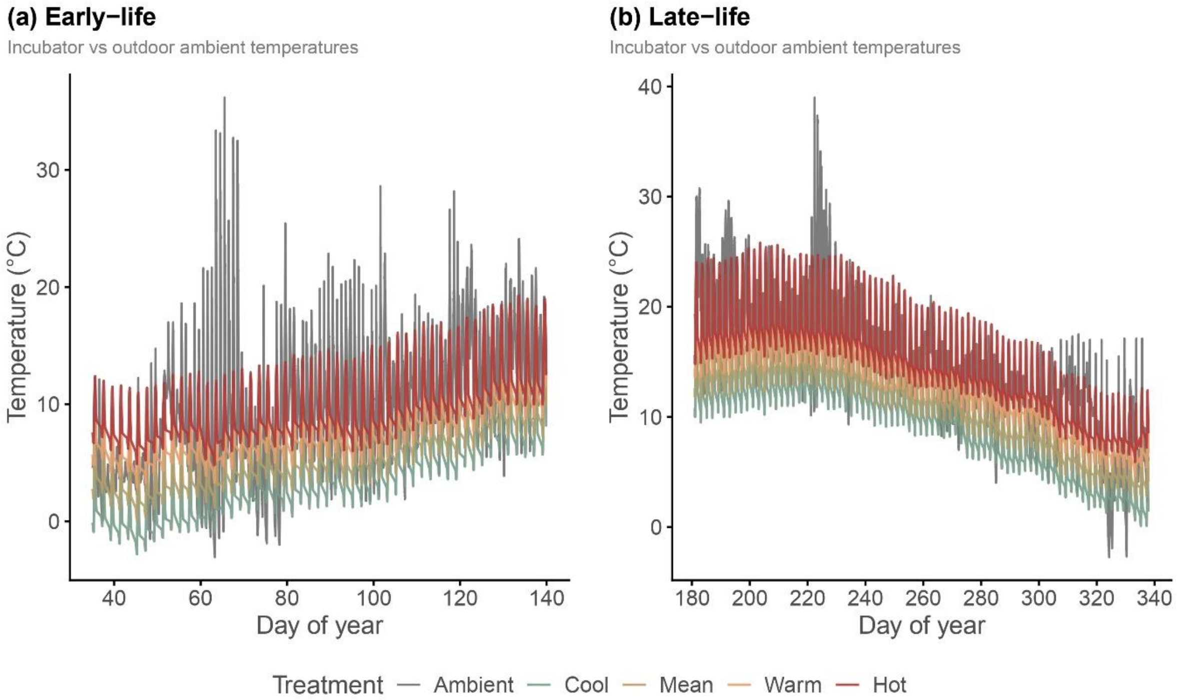


**Figure S3: Temperature profiles for incubator treatments and ambient outdoor treatment. (a)** Temperature from when eggs entered incubators to median timing of pupation. **(b)** Temperature from date pupae entered late-life treatment incubators to median timing of adult emergence.

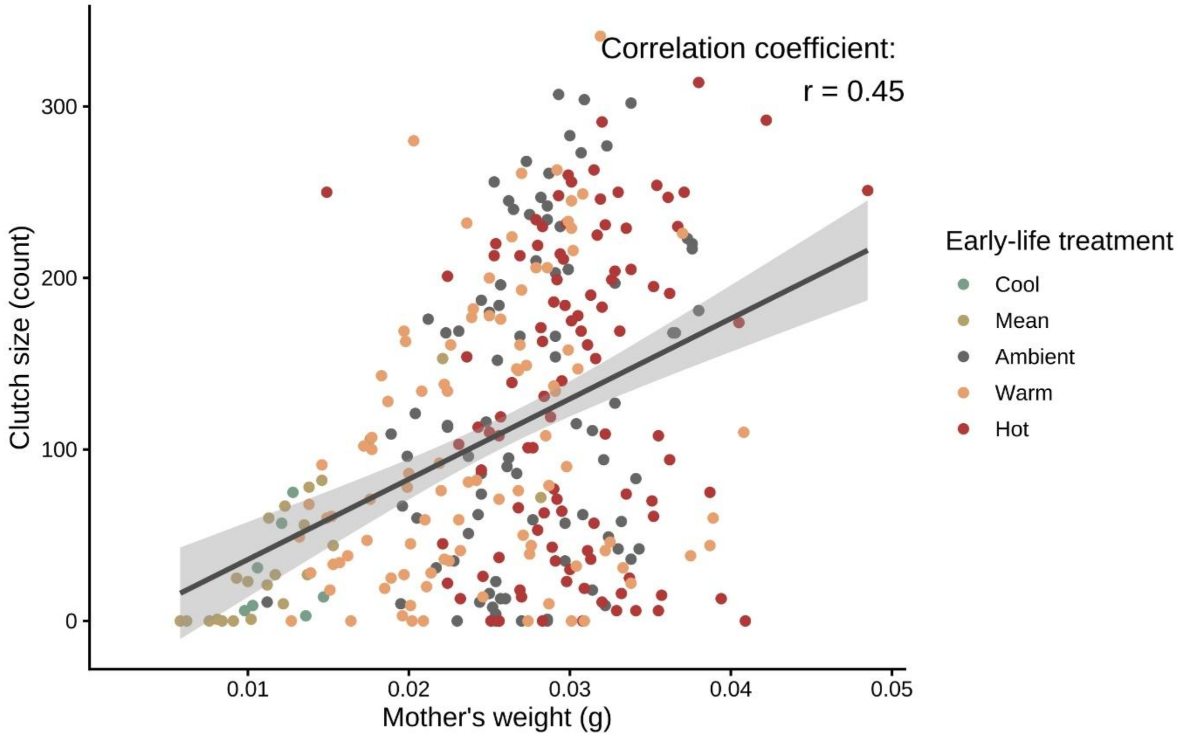


**Figure S4: Mother's weight on adult emergence is correlated with clutch size (r = 0.45).** Figure shows total clutch size per female plotted against her weight on emergence, with points coloured by early-life treatment the mother experienced. Mother's weight is itself associated with early-life treatment experienced, with greater weight associated with warmer early-life treatment.

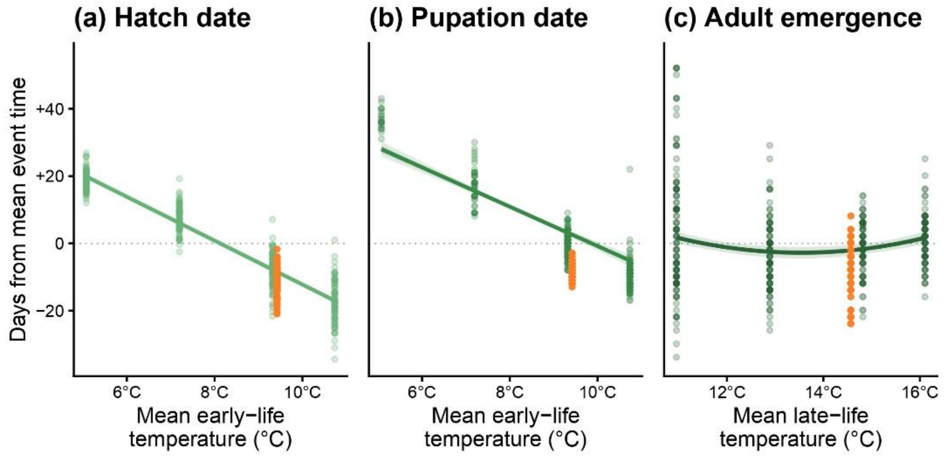


**Figure S5: Ambient group falls along expected line of plasticity to temperature.** Figure is identical to Fig. 2 but shows ambient group raw responses for mean ambient temperature plotted in orange. This shows the treatment groups closely match the expected patterns based on mean temperature at more variable natural conditions. We note that the ambient treatment deviates from the expected pupation-temperature line, possibly due to the influence of occasional days of extreme heat.
